# Neural correlates of individual differences in speech-in-noise performance in a large cohort of cochlear implant users

**DOI:** 10.1101/2021.04.22.440998

**Authors:** Joel I. Berger, Phillip E. Gander, Subong Kim, Adam T. Schwalje, Jihwan Woo, Young-min Na, Ann Holmes, Jean Hong, Camille Dunn, Marlan Hansen, Bruce Gantz, Bob McMurray, Timothy D. Griffiths, Inyong Choi

**Author notes:** Corresponding author at: 250 Hawkins Dr., Iowa City, IA 52242, USA.

## Abstract

**Objectives:** Understanding speech in noise (SiN) is a complex task that recruits multiple cortical subsystems. Individuals vary in their ability to understand SiN. This cannot be explained by simple peripheral hearing profiles, but recent work by our group (Kim et al., 2021, *Neuroimage*) highlighted central neural factors underlying the variance in SiN ability in normal hearing (NH) subjects. The current study examined neural predictors of speech-in-noise ability in a large cohort of cochlear-implant (CI) users, with the long-term goal of developing a simple electrophysiological correlate that could be implemented in clinics.

**Design:** We recorded electroencephalography (EEG) in 114 post-lingually deafened CI users while they completed the California Consonant Test (CCT): a word-in-noise task. In many subjects, data were also collected on two other commonly used clinical measures of speech perception: a word-in-quiet task (Consonant-Nucleus-Consonant [CNC]) word and a sentence-in-noise task (AzBio sentences). Neural activity was assessed at a single vertex electrode (Cz), to maximize generalizability to clinical situations. The N1-P2 complex of event-related potentials (ERPs) at this location were included in multiple linear regression analyses, along with several other demographic and hearing factors as predictors of speech in noise performance.

**Results:** In general, there was a good agreement between the scores on the three speech perception tasks. ERP amplitudes did not predict AzBio performance which was predicted by the duration of device use, low-frequency hearing thresholds, and age. However, ERP amplitudes were strong predictors for performance for both word recognition tasks: the CCT (which was conducted simultaneously with EEG recording), and the CNC (conducted offline). These correlations held even after accounting for known predictors of performance including residual low-frequency hearing thresholds. In CI-users, better performance was predicted by an increased cortical response to the target word, in contrast to previous reports in normal-hearing subjects in whom speech perception ability was accounted for by the ability to suppress noise.

**Conclusions:** These data indicate a neurophysiological correlate of speech-in-noise performance that can be relatively easily captured within the clinic, thereby revealing a richer profile of an individual’s hearing performance than shown by psychoacoustic measures alone. These results also highlight important differences between sentence and word recognition measures of performance and suggest that individual differences in these measures may be underwritten by different mechanisms. Finally, the contrast with prior reports of NH listeners in the same task suggests CI-users performance may be explained by a different weighting of neural processes than NH listeners.

## Introduction

Difficulty understanding speech in background noise (speech-in-noise; SiN) is one of the most common complaints by hearing impaired individuals (for a review, see McCormack & Fortnum, 2013). Indeed, there is considerable variability even across listeners with audiometrically-normal hearing (Fullgrabe et al., 2014; Liberman et al., 2016; Guest et al., 2018). Although peripheral factors may play a role in performance, data from animals and humans suggest that processing within the central auditory hierarchy and beyond contributes significantly to the process of separating signals from noise (e.g. Fullgrabe et al., 2014; Gay et al., 2014; Saiz-Alia et al., 2019).

Previous research suggested that the amplitude of an auditory cortical evoked potential, the N1-P2 complex, reflects the perception of speech at multiple levels (Getz & Toscano, 2020 for a review; Sarrett et al., 2020; Kim et al., 2021). Further, the strength of this component is increased following speech sound discrimination training (Tremblay et al., 2001), demonstrating a clear association between this component and language processing.

Many speech-in-noise tasks elicit two N1-P2 complexes: one to the onset of the noise and one to the onset of the signal. A recent study of speech-in-noise in normal-hearing users capitalized on this to disentangle potential contributors to individual differences in performance. This work demonstrated that SiN performance was well predicted by suppression of the N1-P2 triggered by the onset of the background noise, but not by the N1-P2 to the speech target, nor by activity in higher level frontal areas (Kim et al., 2021). This suggests that, at least in NH listeners, a) early responses to speech may be the most important predictors of performance; and b) the most important contributor to SiN performance is the ability to suppress background noise, rather than boost the target speech.

It is unclear whether this same weighting of processes is evident in hearing-impaired listeners, particularly those using cochlear implants (CIs). Purdy and Kelly (2016) followed 10 CI users for 9 months post-implantation. They showed that N1-P2 responses to non-speech auditory stimuli generally increased, concurrent with significant improvements with speech-in-noise perception ability as listeners adapt to the degraded input of the CI. However, there was considerable variability in evoked responses across subjects, and consequently only P2 amplitudes reached significance. More importantly, tones, rather than speech, were used to elicit the N1-P2, thus it is not clear whether the N1-P2 in this case reflect any processes that are specific to speech.

A number of small-scale cross-sectional studies have also examined various cortical auditory evoked potentials (N1, P2, P300) and their relationship to performance following cochlear implantation. These demonstrate a relationship between the amplitudes of cortical evoked potentials and the speech perception performance in CI users (Micco et al., 1995; Groenen et al., 1996; Makhdoum et al., 1998; Groenen et al., 2001). However, these studies examined cortical responses without background noise, which may utilize only a subset of the cortical mechanisms needed in noise (in support of this point, see Wong et al., 2008; Du et al., 2014). A study by Finke et al (2016) showed a relationship between cortical evoked response to speech in noise and performance in 13 CI listeners. Notably, these studies mentioned above also did not account for other potential predictors of performance that account for variability in understanding speech-in-noise. This can only be done with a large sample.

In order to understand the link between cortical factors and speech-in-noise performance in CI users, it is crucial to set this in the context of the large number of factors that predict how well CI users ultimately adapt following implantation. Many studies show that speech perception improves over time following cochlear implantation (e.g. Tyler et al., 1997; Chang et al., 2010; Blamey et al., 2013; Dillon et al., 2013) but there is large variability across subjects in these gains. A recent systematic review of pre- or peri-lingually deafened CI users summarized several predictors that have been implicated in underlying speech perception performance following late cochlear implantation (Debruyne et al., 2020). These include preoperative speech recognition scores and childhood communication mode. They also noted that there are a number of studies that do show such relationships, but this may be potentially due to low study quality or power.

Many of the largest studies have used large samples of experienced CI users (who have reached asymptotic levels of performance). Gantz et al. (1993) examined a variety of predictors of speech perception in a multiple regression of 48 post-lingually deafened adults. They showed significant influences of duration of profound deafness, speech reading ability, cognitive ability, use of nonverbal communication strategies, engagement in treatment, and degree of residual hearing on speech perception. Similarly, Rubinstein et al. (1999) found that duration of deafness and preoperative sentence discrimination could account for 80% of the variance in performance on a word recognition task, whilst Kitterick and Lucas (2016) also found that shorter duration of deafness and residual hearing in the better ear could account for some of the intra-subject variability in speech perception under difficult listening conditions in CI users.

This work suggests a wide variety of factors that impact performance, though large-scale studies of this kind have not yet examined measures of cortical function such as the N1-P2. The N1-P2 is not likely to be fully independent of these broader factors. For example, duration of deafness affects auditory nerve health; similarly, pre-operative speech perception scores likely reflect both auditory nerve health and residual acoustic hearing that may be available post-implantation. In such cases, these factors likely reflect the quality of the input reaching the cortex, as a critical first determinant of speech-in-noise ability. In that case, the N1-P2 may serve as a biomarker for the summed quality of the peripheral input. This is particularly true for the N1-P2 to the target (not the noise) in a speech-in-noise task, where greater auditory fidelity is required. In contrast, cognitive ability, duration of device use, and engagement in treatment may all affect higher-level cortical adaptation to device use, as a second determinant of speech-in-noise ability. Here, the N1-P2 may serve as a biomarker of global cortical changes derived from multiple causes. In both cases, this predicts that the N1-P2 should predict speech-in-noise performance, but once one accounts for these broader individual factors the N1-P2 should account for little variance.

However, a third possibility is that independently of the hearing loss, individuals vary in the quality of the cortical network for speech-in-noise perception (e.g. Wong et al., 2008; Kim et al., 2021), or word recognition (McMurray et al., 2014). In this case, one might predict an effect of the N1-P2 independent of these other factors. In this way, the N1-P2 could be a useful index of the quality of a listeners’ inherent word recognition and noise suppression abilities.

The current study combined the multifactor approach of the large individual difference studies of outcomes with ERPs to test for a unique link between the N1-P2 (the most promising cortical correlate of performance) and speech-in-noise performance by accounting for the demographic and audiological factors that are known to influence performance. We examined a large cohort of CI users with many histories and device types, with the potential of working towards developing a simple test that could be potentially used in clinics as an objective index of the summed neural response (including both peripheral and central factors) to speech-in-noise.

To this end, we measured neural responses recorded with electroencephalography (EEG) while subjects performed a speech-in-noise task based around the California Consonant Test (CCT; Owens & Schubert, 1977), a closed-set single word task. Performance on this task was used as a dependent variable, along with two other speech perception tasks used in clinical evaluation, conducted in separate sessions (without EEG) and using different testing formats: an open set speech-in-quiet task (consonant-nucleus-consonant [CNC]; Lehiste & Peterson, 1959), and a sentence-in-noise task (AzBio; Spahr et al., 2012).

Performance on these three dependent variables was related to the N1-P2 complex in the EEG as well as other relevant audiological and demographic factors. Unlike prior studies, we focused on a large population of hybrid and bimodal CI users who combine acoustic hearing with residual low frequency acoustic hearing, either in the ear contralateral to the CI (a bimodal configuration) or the ipsilateral ear (hybrid). This allowed us to include low frequency acoustic hearing thresholds. Given that this is nearly always beneficial to word recognition (Gifford et al., 2013), this was expected to be a strong marker of a strong auditory periphery. We used a relatively simple ERP component – the N1-P2 which can be measured from a single electrode (Cz), rather than more complex ICA or source analysis techniques. This was done for two reasons. First, the N1-P2 - even at this single electrode - has been robustly related to language and speech perception in noise (Tremblay et al., 2001; Sarrett et al., 2020; Kim et al., 2021). Second, our hope was that a simpler measure may prove more useful diagnostically in clinical settings. The long-term goal here is that by understanding both cortical and demographic factors underlying variance in speech perception performance across hearing impaired listeners for different tasks, this could help to inform and refine remediation strategies.

## Materials and Methods

### Participants

One hundred and fourteen CI users, between 18 and 85 years of age (mean = 62.6 years, SD = 13.5 years; median = 65.8 years; 59 (51.8%) female), were recruited from the University of Iowa Cochlear Implant Research Center. All the participants were post-lingually deafened and neurologically normal. Average length of device use was 39.5 months (SD = 56.8 months). Average duration of deafness was 22.0 years (SD = 15.0 years). Five subjects were bilateral CI users. Among the remaining one hundred and nine subjects, seventy-two subjects (66.1%) had CI in the right ear. Eighty-seven subjects (76.3%) were Hybrid CI users (i.e., electric acoustic stimulation within the same ear). Average threshold of low-frequency (i.e., 250 and 500 Hz) residual acoustic hearing was 59.4 dB HL (SD = 20.5 dB HL). American English was the primary language for all the participants. Most participants were tested during the same day as a clinical visit in which they received an annual audiological examination and device tuning. Full details of all participants (e.g. demographics, device types) can be found in *Supplementary Table 1*. Critically, most of these CI users were bimodal or hybrid CI users who would have some residual acoustic hearing. Duration of device use was obtained from clinical records. All study procedures were reviewed and approved by the local Institutional Review Board.

### Task design and procedures

All CI users performed the CCT simultaneously as EEG was recorded. In a subset of users, we also obtained performance on two commonly-used clinical tests for speech perception: CNC word recognition (in quiet) (*n* = 89) and AzBio sentence recognition (in noise) (*n* = 72). These were conducted by a trained audiologist in a separate session, usually on the same day. CNC and AzBio scores were collected within the clinic during routine audiological examinations. Subjects were excluded from these sub-analyses if we did not have complete data, hence the smaller numbers of subjects for these other two tasks.

#### CCT

The CCT was implemented in custom-written Matlab scripts (R2016b, Mathworks) using the Psychtoolbox 3 toolbox (Brainard, 1997; Pelli, 1997). The CCT (and the simultaneous EEG recording) was conducted in an acoustically-treated, electrically-shielded booth with a single loudspeaker (model LOFT40, JBL) positioned at a 0° azimuth angle at a distance of 1.2 m. Visual stimuli were presented via a computer monitor located 0.5m in front of the subject at eye level. Sound levels were the same across subjects.

The overall structure of the trials for the CCT/EEG paradigm is shown in Figure 1. For each trial, participants saw a fixation cross on the computer screen. They were instructed to fix their gaze on the cross for the duration of the trial, to minimize eye-movement artifacts on the EEG. They were then presented with an auditory cue phrase (“check the word”, ~800 ms duration, spoken by the same talker as target words), to prime them for the onset of the background babble. Following a 700 ms period of silence, eight-talker babble (herein referred to as the “noise”) began and continued for 2 seconds. Target words were presented 1 second after noise onset. Target words were always presented at 70 dB SPL and consisted of 100 monosyllabic consonant-vowel-consonant words selected from the CCT, spoken by an American male talker with a Midwest accent. The noise was varied randomly between two levels: +7 dB (low signal-to-noise ratio; SNR) and +13 dB (high SNR), relative to the target word. For each SNR, 50 trials were presented. The division of 100 words into two sets of 50 words was fixed across listeners, although the presenting order was randomized. The phonetic balance in each SNR condition is described in Supplementary Table 2 (used from Kim et al., 2021). Finally, 100 ms following the offset of the target acoustic stimulus, participants were presented with a four-alternative forced choice task, seeing four possible words printed in the center of the screen, stacked vertically and labeled with a corresponding number. They then used a numeric keypad to select the target word they thought they had heard. No feedback was given regarding correctness and trials did not end until 1 second after a response was given.

**Figure 1.**
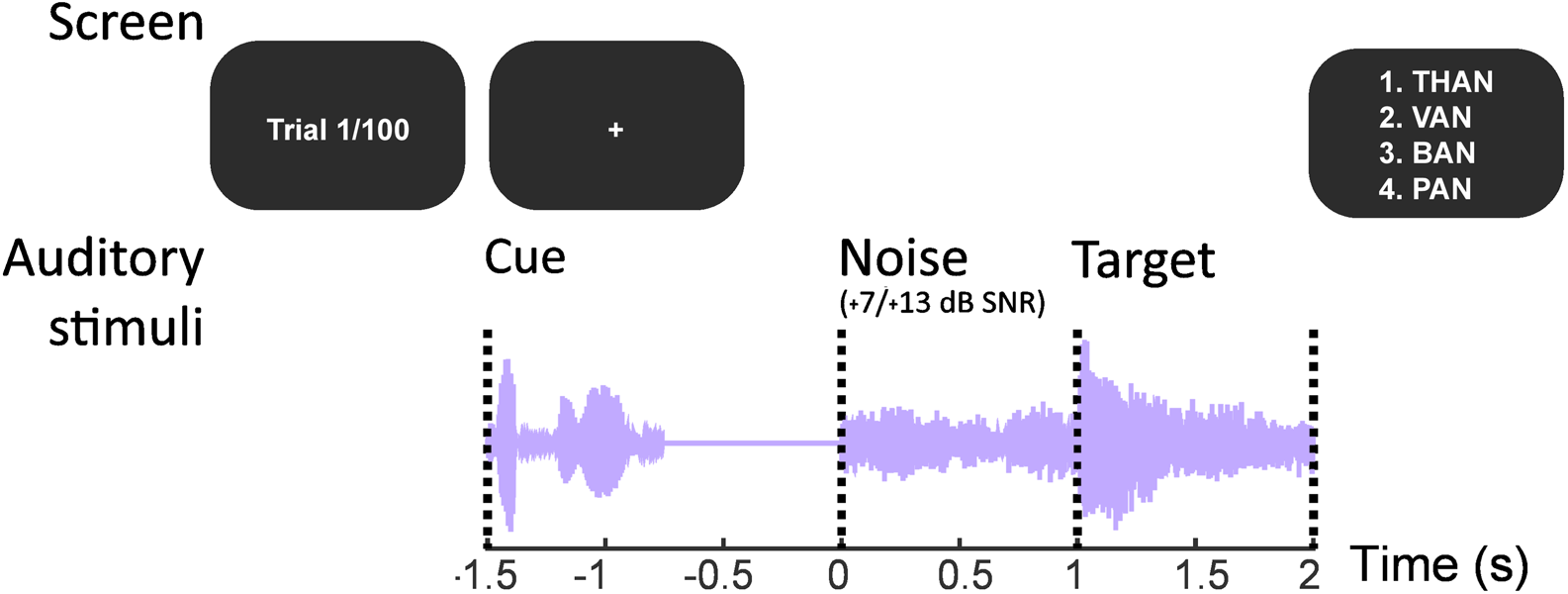
Trial and stimulus structure for CCT paradigm. Each trial began with a fixation cross, followed immediately by the cue phrase (“check the word”). Background noise onset began 700 ms following cue offset and target words began 1 second after this. Subjects were then required to respond to a 4-choice alternative task using a 4-button keypad. The noise level was manipulated relative to the target word, to create either high (+13 dB) or low (+7 dB) SNR conditions. A total of 50 trials was presented for each condition.

### EEG acquisition and pre-processing steps

EEG data were recorded using a BioSemi ActiveTwo system at a 2048 Hz sampling rate, with either 64 or 128 electrodes arranged in a 10-20 placement system. Physiological data and stimulus-timing triggers were recorded in ActiView (BioSemi). Data from each channel were filtered offline between 1 to 50 Hz with a 2048-point FIR filter. We then epoched data around the stimulus from 0.5 s before the noise onset to the appearance of the response options. Next, sound-evoked artifacts introduced by CIs and eye-blink artifacts were removed using independent component analysis (implemented in the Matlab EEGLab toolbox; Delorme & Makeig, 2004). CI artifact-related components were determined by the combination of visual inspection and an automated process that observed temporal and spatial patterns. Eye blink artifacts were determined by visual inspection. Baseline correction was applied by subtracting the mean voltage between −200 and 0 ms prior to noise onset, and epoched data were down-sampled to 256 Hz.

Although we recorded from up to 128 electrodes, these data were used solely for the purposes of visualizing the topography of cortical responses (see Figure 2). Our long-term goal was to consider the utility of this method for clinical purposes, where it may only be feasible to collect a limited number of electrodes. Thus, only the Cz electrode was included in the subsequent statistical analyses. All ERPs were z-scored prior to further analyses to reference by common averaging. N1-P2 amplitudes were extracted by determining the maximal negative and positive waveforms within a 400 ms window following the onset of a stimulus. We then subtracted the N1 from the P2 to obtain a composite measure. This was done separately for the N1-P2 triggered by the noise onset, and the N1-P2 triggered by the target speech.

**Figure 2.**
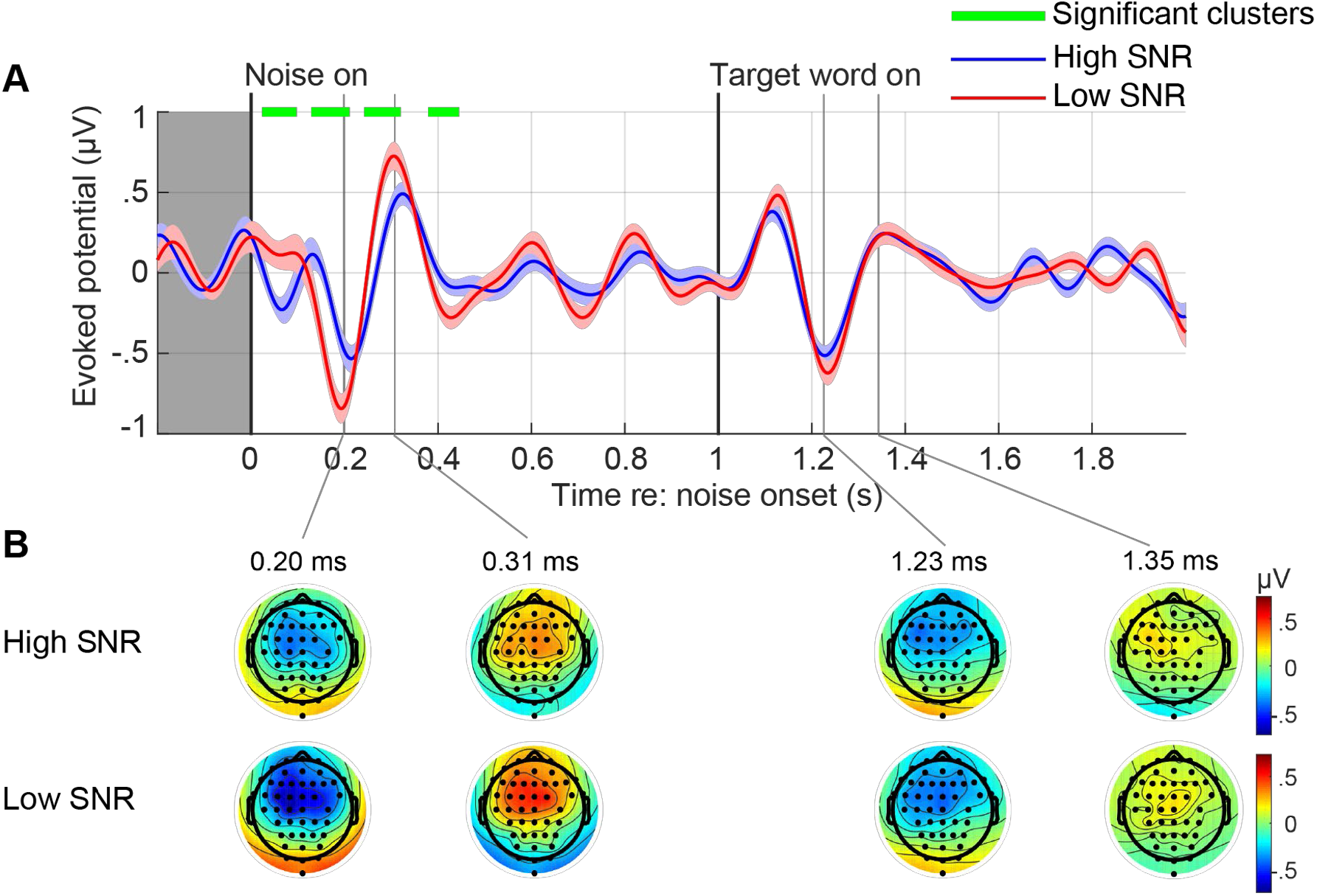
A: Grand-average ERPs (± standard error; *n* = 114) at electrode location Cz for each condition are displayed. Green bars indicate significant clusters (cluster-based permutation test). B. Topographies are plotted for the N1 and P2 responses to both the noise (first two columns) and the target (final two columns), for both conditions (top and bottom rows).

### Statistical analyses

The ultimate goal of this project was to relate speech perception performance to ERP components during speech in noise processing, while controlling for known demographic and audiological factors. For this we employed three multiple regression analyses. Each predicted accuracy on one of the three speech perception measures (CCT [N=114], CNC [N=89], and AzBio [N=72]). Our plan was to first examine the data with bivariate analyses, to examine the spread of the data and assess other factors, such as collinearity. We then conducted large-scale multivariable analyses, predicting each performance outcome from ERP measures and individual factors

We considered two ERP measures as potential predictors of performance: the N1-P2 amplitudes to the target word (N1-P2_target_), and the N1-P2 amplitudes to the noise (N1-P2_noise_). The relationship of the EEG measures to performance was assessed accounting for several other factors. These included duration of device use (log-scaled in months; DeviceDuration), average of residual low frequency hearing thresholds (250 and 500 Hz; PTA_low_) and current age (Age).

We then related each predictor to speech perception performance on 1) CCT task (which was conducted simultaneously with the EEG), CNC task or AzBio, in three separate multiple regression models. The final model is given in (1), in the syntax of the lm() function in R. For the CCT task, the dependent variable (Speech Perception) was the mean accuracy in the low SNR condition. This condition was selected as it is the most difficult and therefore best reflects a subject’s difficulty in understanding speech-in-noise. For the CNC and AzBio tasks, mean performance on each task was used as a dependent variable in separate models.

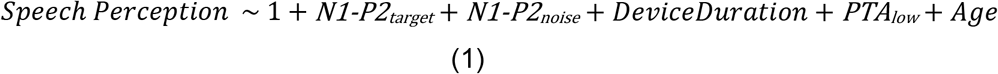

The regressions were limited to the specific N1-P2 ERP components for the reasons described above. A key factor when in a multiple linear regression is the normality of the residuals. Normality testing was also performed on the residuals for each dependent variable using Shapiro-Wilk tests. All variables tested were confirmed to meet this prerequisite.

We also conducted exploratory analyses on the full time-course of the EEG signal, by collapsing listeners into two categories (good and poor performers) - defined using a median split - and comparing them at each timepoint using a cluster-based permutation test (Maris & Oostenveld, 2007). Significant clusters were included if they lasted for a minimum of 50 ms (i.e., 13 samples at 256 Hz sampling rate). A similar approach was conducted by collapsing trials in the CCT into the high and low SNR conditions. All statistical analyses were carried out using custom-written Matlab scripts.

## Results

### Performance on the three different tasks

Mean performance accuracy across subjects (± s.d.) is shown in Supplementary Figure 1. Performance was similar for both of the speech-in-noise tasks (single-word CCT = 52.95% ± 15.69; sentence-based AzBio = 50.93% ± 26.56), whilst the best performance was on the word-in-quiet task (CNC = 68.85% ± 17.99).

### CCT ERP analyses

#### Effect of SNR on ERP

We first examined the effect of SNR on the ERP amplitudes to verify data quality and examine topographical maps of the response. Figure 2 shows the ERP at electrode Cz to the noise and the target on the simultaneous CCT task for both the low and high SNR conditions, averaged across subjects (*n* = 114). A clear N1 and P2 can be seen approximately 200 ms after noise onset, and again about 200 ms after the target word onset. Topographic maps at components peaks (the N1 and P2 to the noise and target) demonstrated that, as expected, peak amplitudes were located at or near location Cz.

We observed an enhanced ERP response to the noise in the low SNR condition compared to the high SNR condition. This appeared to span both the hypothesized N1-P2 components, but also the earlier points (the P1 or P50). However, this was not seen in the N1-P2 to the target. This is an unsurprising result, as the noise stimulus was increased by 6 dB in this condition, whereas the target stimulus level was kept fixed.

#### Median split of good vs poor performers

Work from our own group recently demonstrated that ERP responses to the noise in a CCT task with NH listeners could distinguish good from poor performers (Kim et al., 2021). Thus, we next sought to address the same question in CI users. To this end, we plotted grand averaged ERPs to the low SNR condition at location Cz following a median split of CCT performance (Figure 3). The clearest and significant differences – assessed with a cluster-based permutation analysis - were present at the timing of the N1 and P2 to the target. Topographic maps again confirmed that these effects were pronounced around location Cz.

**Figure 3.**
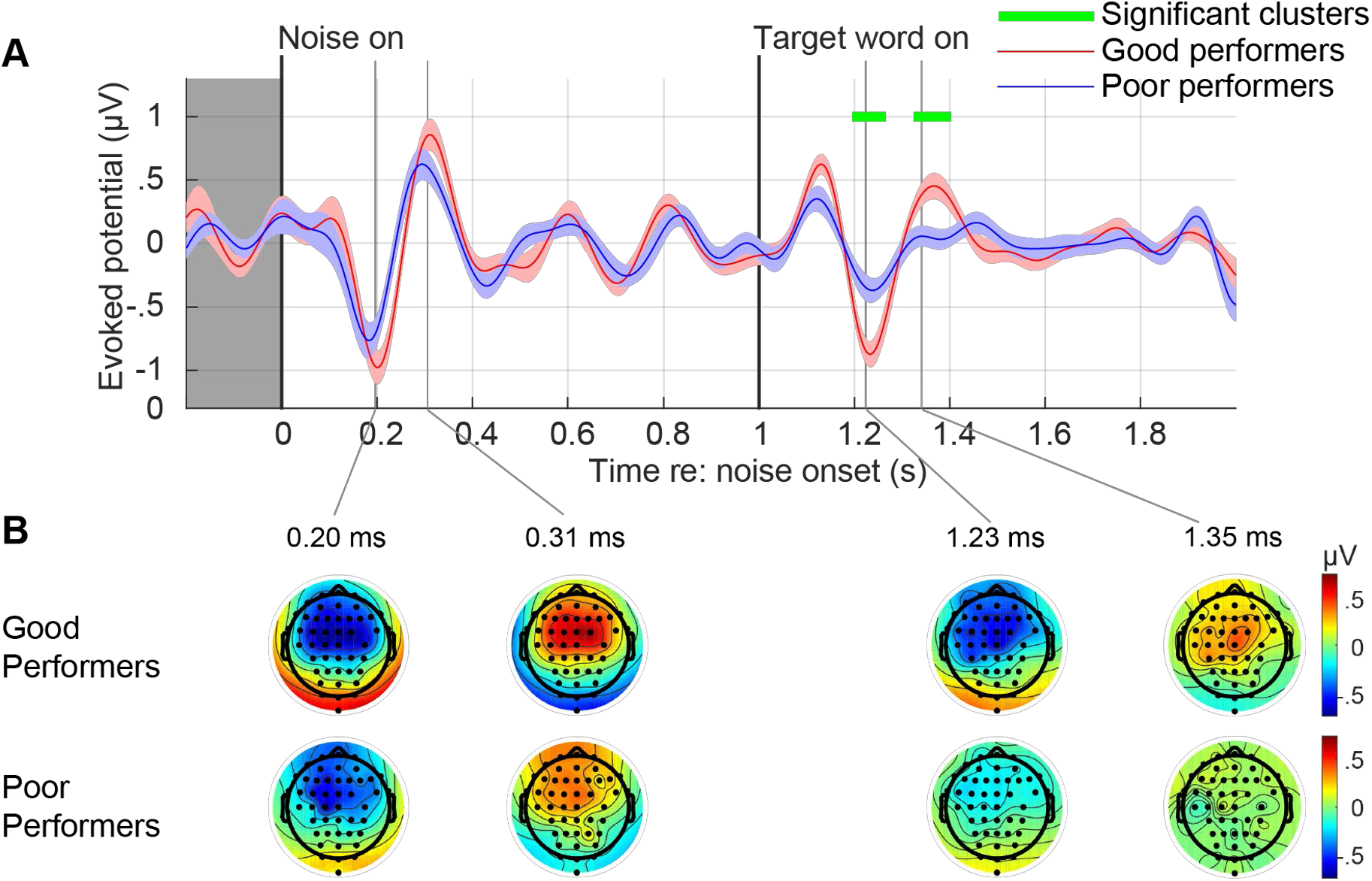
A: Median-split ERP comparisons of good performers (*n* = 57, blue line) vs. poor performers (*n* = 57, red line), in response to the low SNR condition on the CCT task. Shading indicates standard error. Green bars indicate significant clusters. B: Amplitude topographies for good performers and poor performers, for the N1 and P2 responses to the noise and target, are shown below the line plot.

### CCT predictive analyses

#### Evaluation of independent variables in bivariate analyses

We started by evaluating the correlations among all the independent variables, in order to check for co-linearity prior to multiple linear regression analysis (see Supplementary Figure 2). None of the variables reached the threshold for being considered colinear (|r| ≥ 0.7; Dormann et al., 2013). Nonetheless, there was a small but significant correlation between duration of implantation (DeviceDuration) and residual low frequency hearing threshold (PTA_low_; r = 0.21, *p* = 0.025), suggesting that low frequency acoustic hearing worsens over the duration of device use. Although the majority of devices included here were a hybrid design, which can greatly preserve low frequency hearing, some degree of loss over time has been shown previously (Gantz et al., 2009). There was also a small but significant negative correlation between N1-P2_target_ and Age (r = −0.22, *p* = 0.020) with a reduction of the ERP amplitudes in older listeners (Figure 4B, left panel). These correlations suggest that while no variables are collinear, they are not unrelated, underscoring the need for a multiple regression.

**Figure 4.**
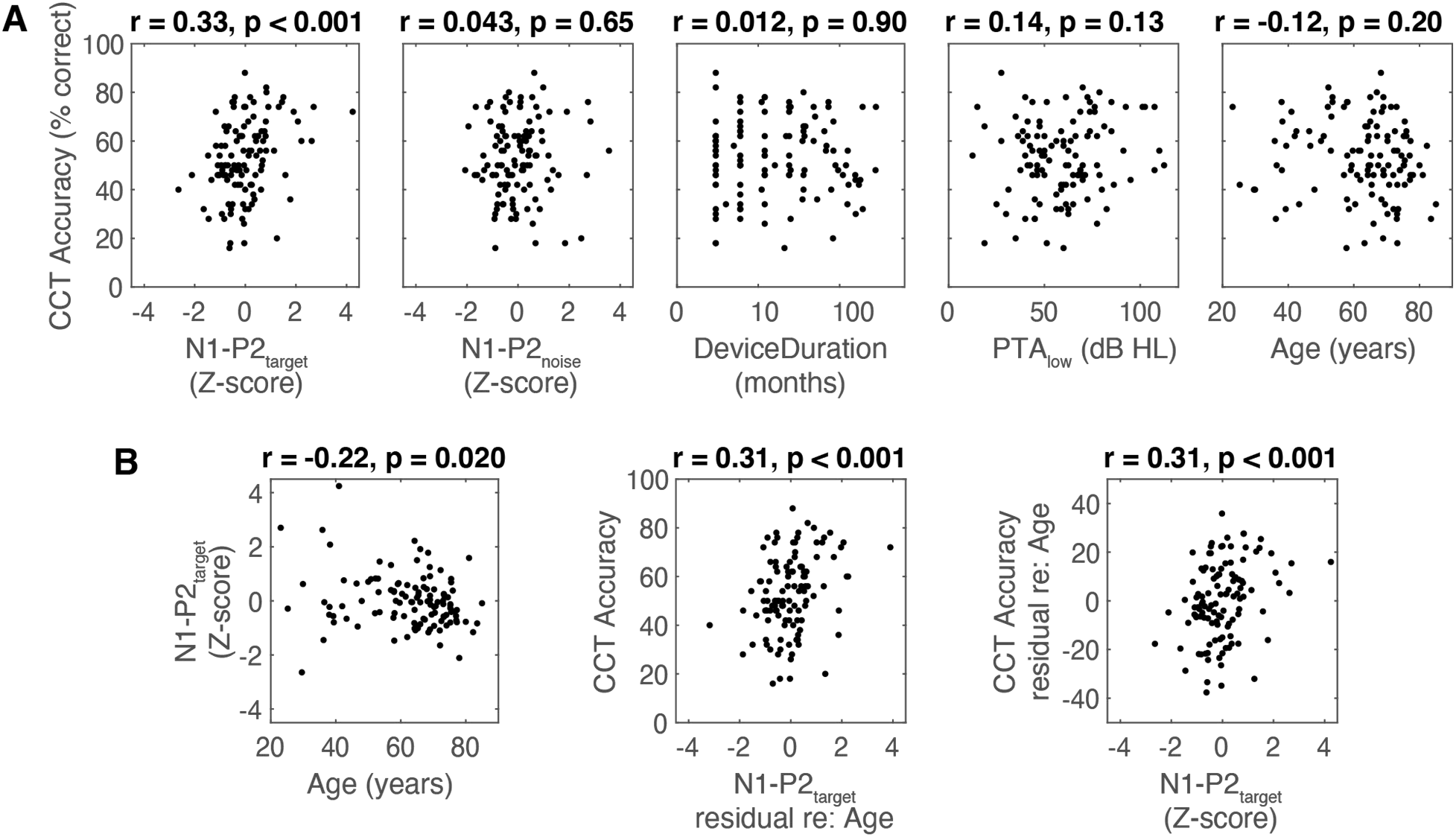
Results from bivariate analyses. A. Each predictor plotted against CCT accuracy, with r and *p* values displayed from bivariate analyses (*n* = 114). B. Regression of N1-P2 amplitudes against age (left panel), as well as bivariate analyses following residualization of either N1-P2 against age (middle panel) or CCT accuracy against age (right panel).

Next, we conducted exploratory bivariate analyses examining correlations between each independent variable (both the ERP and non-ERP variables) and CCT accuracy in the low SNR condition. These are shown in Figure 4A (and see Supplementary Figure 2). The only significant correlation was a positive one between N1-P2_target_ and CCT accuracy (r = 0.33, *p* < 0.001), with larger amplitudes predicting better performance in the low SNR condition. Other predictor variables did not exhibit significant correlations.

Since age had an effect on the amplitude of the N1-P2_target_, we sought to isolate an effect of the N1-P2 that was independent of age. We thus, residualized the N1-P2_target_ against age and examined the correlation between the residualized N1-P2 amplitudes to the target and accuracy on the CCT. These results are shown in Figure 4B. N1-P2 target amplitudes were still significantly correlated with CCT accuracy after regressing out the effect of age (r = 0.31, *p* < 0.001). This was also true when age was residualized out of CCT accuracy (r = 0.31, *p* < 0.001).

#### Multiple linear regression - CCT_low_ as dependent variable

Following bivariate analyses, we conducted a multiple linear regression analysis to determine which of the independent variables predicted CCT accuracy when accounting for all others (see Table 1 in its entirety). When adjusted for the number of independent variables, the model accounted for 13.5% of the variance in CCT accuracy, *F*(5, 108) = 3.38, *p* = 0.007, Adjusted *R^2^* = 0.095. The only significant predictor was amplitude of the N1-P2 to the target, *F*(1, 112) = 12.1, *p* < 0.001. Given the presence of age, PTA_low_ and length of device use in the model, this confirms an effect of the EEG even after accounting for this. It is also notable that, unlike in NH listeners (Kim et al., 2021), N1-P2 to the *noise* was not a significant predictor of performance, *F*(1, 112) = 0.015, *p* = 0.90.

**Table 1:** Results from multiple linear regression on CCT accuracy (N=114, R^2^=0.14):

| **CCT<sub>low</sub>** | **$\beta$** | **SE** | **T(112)** | **$p$** | **Partial $\rho$** |
| --- | --- | --- | --- | --- | --- |
| **N1-P2<sub>target</sub>** | 5.11 | 1.47 | 3.5 | **0.00071** | **0.32** |
| **N1-P2<sub>noise</sub>** | 0.18 | 1.45 | 0.12 | 0.90 | 0.012 |
| **DeviceDuration** | -0.78 | 2.44 | -0.32 | 0.75 | -0.031 |
| **PTA<sub>low</sub>** | 0.12 | 0.071 | 1.7 | 0.10 | 0.16 |
| **Age** | -0.052 | 0.11 | -0.48 | 0.63 | -0.046 |

Next, to determine the generalizability of these effects, we further examined the predictors against performance data collected on the other two speech perception tasks.

### CNC

#### Evaluation of independent variables in bivariate analyses

As these data included only a subset of the CI users (those for which we also had either CNC or AzBio behavioral data), we again examined the predictor variables in bivariate analyses for the CNC task. As with the CCT, none of the independent variables reached the threshold for collinearity for either analysis. In this subset of users, there was still a significant correlation between DeviceDuration and PTA_low_ (r = 0.30, uncorrected *p* = 0.005). There was also still a significant negative correlation between N1P2_target_ and Age (r = −0.34, uncorrected *p =* 0.001). Results from bivariate analyses examining correlations between each independent variable and CNC accuracy are shown in Figure 5A. Similar to the CCT, there was a significant positive correlation between N1P2_target_ and CNC accuracy (r = 0.32, uncorrected *p* = 0.002). There was also a significant correlation between DeviceDuration and CNC accuracy (r = 0.25, uncorrected *p* = 0.018), and Age and CNC accuracy (r= −0.23, uncorrected *p* = 0.028). None of the other variables correlated significantly with accuracy.

**Figure 5.**
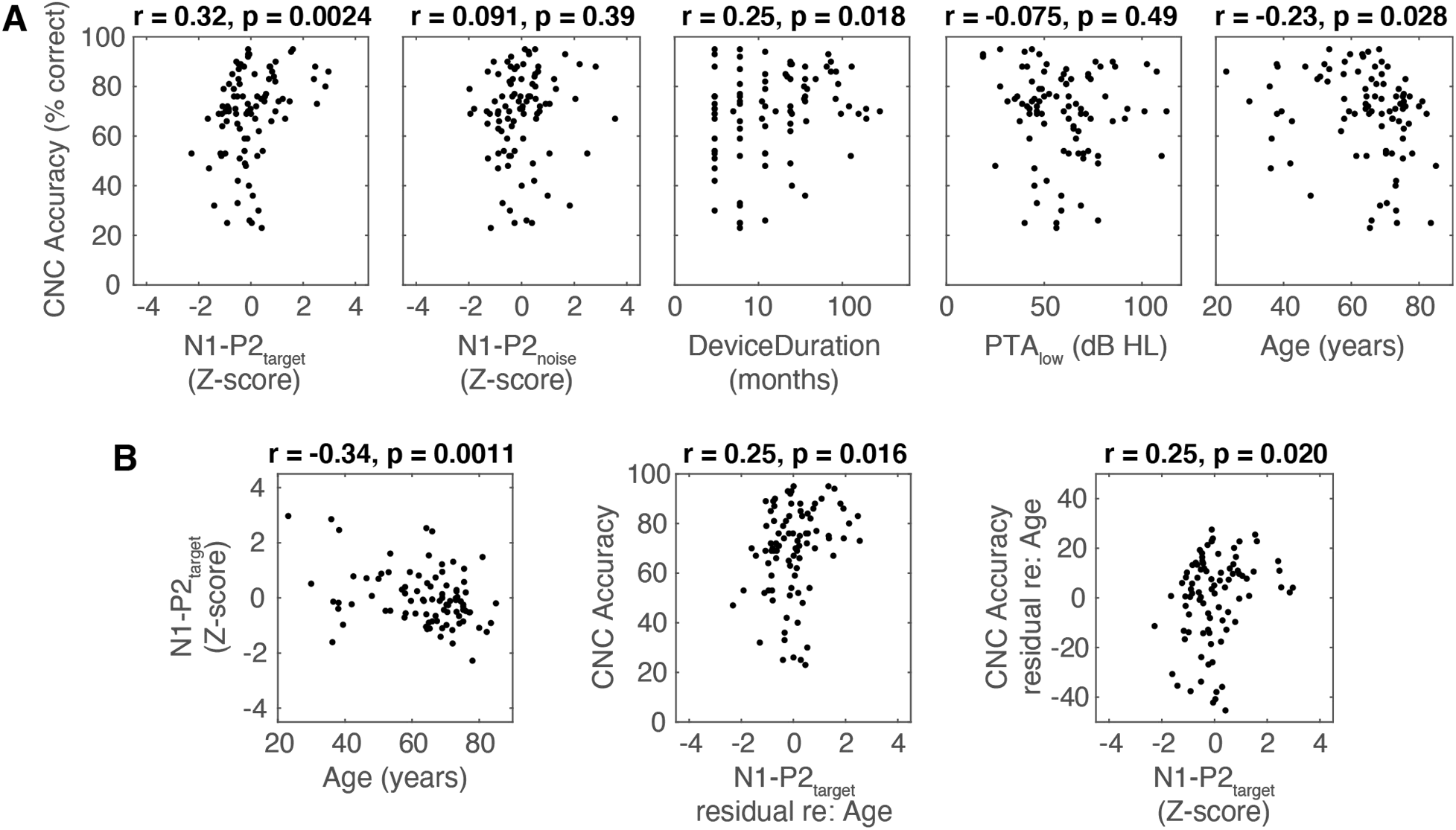
Predicting CNC accuracy. A: Predicting accuracy on the CNC task with bivariate analyses (n = 89). B. Regression of N1-P2 amplitudes against age (left panel), as well as bivariate analyses following residualization of either N1-P2 against age (middle panel) or CNC accuracy against age (right panel).

Again, as with the CCT dataset, the significant correlation between N1-P2_target_ and CNC accuracy persisted even following residualization against age (r = 0.25, *p* = 0.016; Figure 5B).

#### Multiple linear regression – CNC accuracy as dependent variable

The results of a multiple linear regression analysis – with CNC accuracy as the dependent variable – are displayed in Table 2. Following adjustment for the number of independent variables, this model accounted for 19.3% of the variance in CNC accuracy, F(5, 83) = 4.0, *p* = 0.003, Adjusted *R^2^* = 0.15. As with the CCT accuracy, the amplitude of the N1-P2 to the target measured during the CCT test was a significant predictor of CNC accuracy, *F*(1, 87) = 5.9, *p* = 0.018. Additionally, duration of implantation was a predictor of CNC accuracy, *F*(1, 87) = 6.8, *p* = 0.011.

**Table 2:**
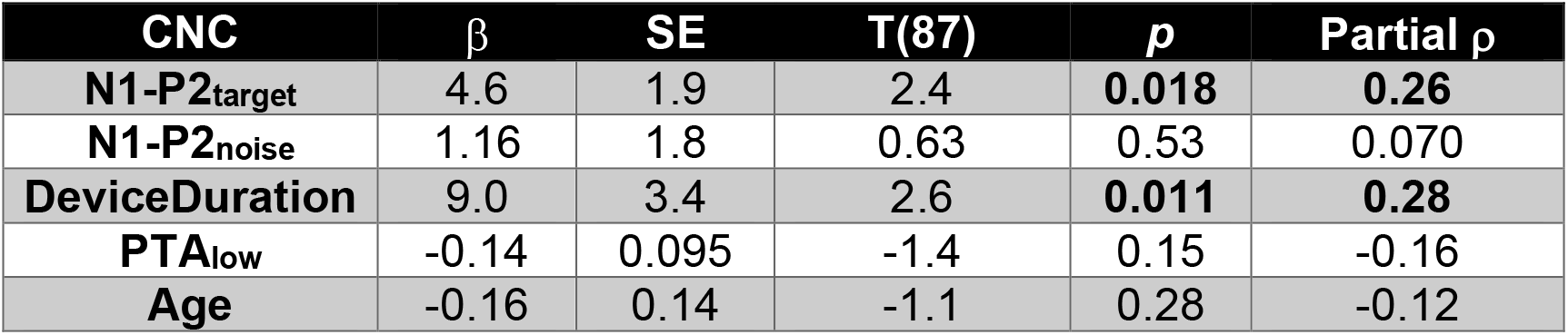
Results from multiple linear regression on CNC accuracy (N=89, R^2^=0.19):

| CNC | $\beta$ | SE | T(87) | $p$ | Partial $\rho$ |
| --- | --- | --- | --- | --- | --- |
| **N1-P2<sub>target</sub>** | 4.6 | 1.9 | 2.4 | **0.018** | **0.26** |
| **N1-P2<sub>noise</sub>** | 1.16 | 1.8 | 0.63 | 0.53 | 0.070 |
| **DeviceDuration** | 9.0 | 3.4 | 2.6 | **0.011** | **0.28** |
| **PTA<sub>low</sub>** | -0.14 | 0.095 | -1.4 | 0.15 | -0.16 |
| **Age** | -0.16 | 0.14 | -1.1 | 0.28 | -0.12 |

### AzBio task predictive analyses

#### Evaluation of independent variables in bivariate analyses

Finally, we turned to the AzBio sentence recognition task (*n* = 72). Predictor variables were again first examined with bivariate analyses prior to implementing multiple linear regression. The only significant correlation between independent variables for this subset of users was between N1-P2_target_ and Age (r = −0.30, *p* = 0.012).

Figure 6A shows the same analyses applied to AzBio accuracy. Unlike the other two tasks, N1P2 amplitudes did not significantly correlate with AzBio accuracy (r = 0.13, *p* = 0.27). Furthermore, residualization against age did not reveal any significant correlation (Figure 6B). In the bivariate analyses, age was negatively correlated with AzBio accuracy (r = −0.31, *p* = 0.008).

**Figure 6A:**
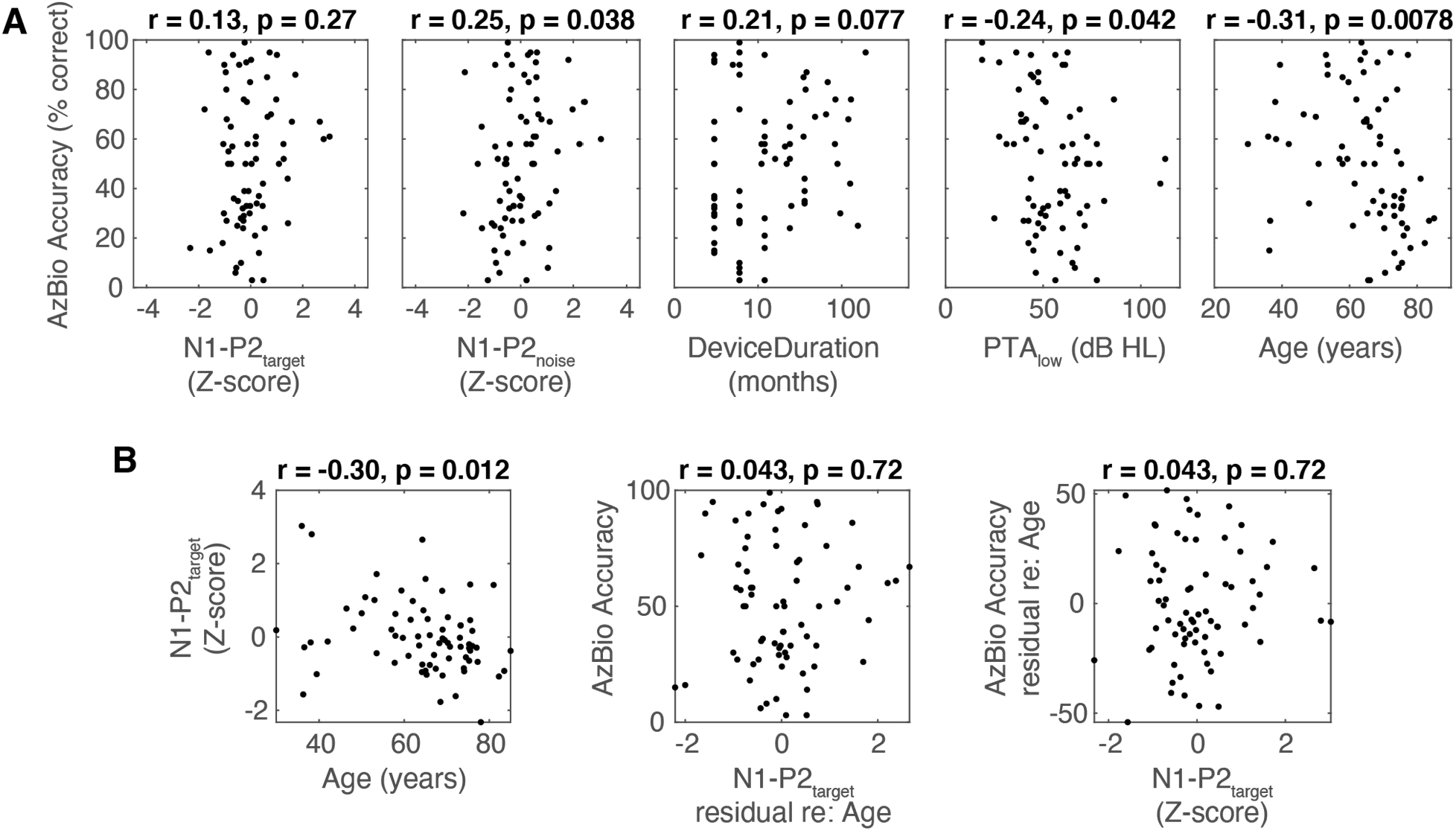
Predicting accuracy on the AzBio task with bivariate analyses (n = 72). B. Regression of N1-P2 amplitudes against age (left panel), as well as bivariate analyses following residualization of either N1-P2 against age (middle panel) or AzBio accuracy against age (right panel).

#### Multiple linear regression – AzBio accuracy as dependent variable

Table 3 shows the results of a multiple linear regression with AzBio accuracy as the dependent variable. Overall, the model accounted for 22.1% of the variance in AzBio accuracy, *F*(5, 66) = 3.75, *p* = 0.005. Interestingly, unlike in the other tasks, the N1-P2 to the target was not significantly correlated with AzBio performance (*F*(1, 70) = 0.012, *p* = 0.91). However, a number of the other individual factors did reach significance. These included the duration of device use (DeviceDuration; *F*(1, 70) = 4.4, *p* = 0.039), the amount of residual acoustic hearing (PTA_low_; *F*(1, 70) = 4.7, *p* = 0.034) and age (Age; *F*(1, 70) = 4.8, *p* = 0.032).

**Table 3:**
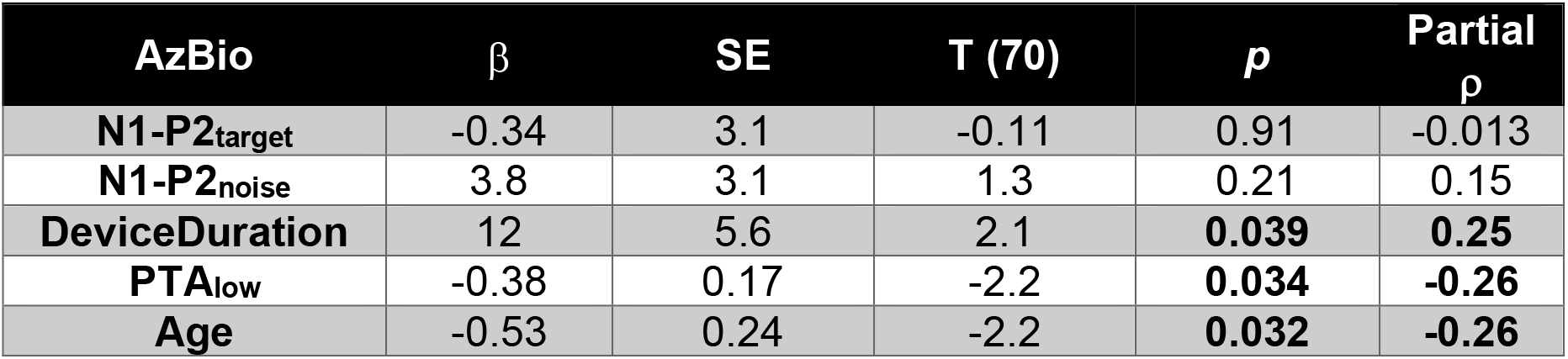
Results from multiple linear regression on AzBio accuracy (N=72, R^2^=0.22):

| AzBio | $\beta$ | SE | T (70) | $p$ | Partial $\rho$ |
| --- | --- | --- | --- | --- | --- |
| **N1-P2<sub>target</sub>** | -0.34 | 3.1 | -0.11 | 0.91 | -0.013 |
| **N1-P2<sub>noise</sub>** | 3.8 | 3.1 | 1.3 | 0.21 | 0.15 |
| **DeviceDuration** | 12 | 5.6 | 2.1 | **0.039** | **0.25** |
| **PTA<sub>low</sub>** | -0.38 | 0.17 | -2.2 | **0.034** | **-0.26** |
| **Age** | -0.53 | 0.24 | -2.2 | **0.032** | **-0.26** |

## Discussion

The current study demonstrated that the amplitude of the N1-P2 response to target words was uniquely related to performance on single word identification tasks in CI users, even after accounting for other potentially confounding demographic factors. This was shown in our primary measure (the CCT). But it also generalized to a quite different speech perception task (open set), using different stimuli (a different talker and different words), in quiet, and conducted in a separate session. This provides robust evidence for a link between the N1-P2 and single word recognition. Crucially, the single-word results during the EEG task confirm one of the hypotheses described in the introduction - that is, independent of other likely predictors of speech perception (e.g. residual low frequency hearing preservation), individuals vary in the quality of their cortical network for speech-in-noise perception.

Interestingly, whilst the duration of device use, residual low frequency hearing, and age were related to performance on a sentence-in-noise task (AzBio), the N1-P2 response was not. We should note that there were fewer data points contributing to the AzBio analyses, although we had *n* = 72, so this should still be sufficient to detect an effect (Harris, 2001). However, looking across both tasks, we see a substantially different profile of predictors, with single word tasks being largely related to auditory-cortical factors, and sentence tasks reflecting a range of other factors. These results highlight important differences between the single-word and sentence-based tasks, which we will discuss below.

### Demographic predictors of speech perception performance

It is intriguing that most of the usual demographic factors did not predict single word performance in either the CCT or the CNC, both individually and in the multiple regression. These measures of SiN performance were only related to the N1-P2 (although duration of device use was also a predictor for the CNC). Based on previous studies presented here, this is at first a surprising result. In contrast, the clearest predictors of AzBio behavioral performance were the duration of device use, the degree of residual low frequency hearing threshold, and age. This highlights the importance of preserving low frequency hearing when possible and is consistent with prior observations that preservation of low frequency hearing significantly improves speech perception and music appreciation (Turner et al., 2004; Gantz et al., 2005; Dunn et al., 2010).

It is nonetheless surprising that performance on the single word paradigms was not predicted by residual low frequency thresholds in the current study. This should be investigated further, given the evidence from previous studies mentioned above indicating a benefit of residual low frequency hearing. However, we note that the large majority of previous studies use within-subject manipulations to study the benefit of acoustic hearing (e.g. comparing performance with the CI only to performance with the CI and hearing aid) – there are fewer studies that evaluate the benefit of acoustic plus electric (A + E) hearing between subjects as was the case here (e.g. Green et al., 2007; Gifford et al., 2013; Kitterick & Lucas, 2016). It is also important to note that for most of these patients (as is often the case with A+E hearing), the majority of acoustic hearing comes from low frequencies. These frequencies may contribute little to single word recognition (since most phonetic cues necessary for contrasting words are above 500 Hz). In contrast, in sentences these frequency bands could help with word segmentation and with grouping or streaming the target sentence from the background noise. Furthermore, the benefits of acoustic hearing must be considered more broadly than just speech-in-noise (e.g. talker identification, musical appreciation); therefore, these results do not mitigate the importance of preserving low frequency hearing thresholds during cochlear implantation, as this undoubtedly still has important quality-of-life benefits. Moreover, the ecological validity afforded by the sentence-in-noise paradigms (Taylor, 2003) further supports the idea that preservation of low frequency hearing thresholds likely provides benefits in common real-world listening scenarios.

### N1-P2 amplitudes in NH and CI users

In our recent study of normal hearing listeners (Kim et al., 2021), speech-in-noise performance was best predicted by suppression of the N1-P2 response to the noise, rather than enhancement of the N1-P2 to the target. This contrasts with the finding here that increased N1-P2 amplitudes to the target indicated better single word speech perception across CI users.

There are two main possible explanations for this. One is that CI users are using different cortical mechanisms for speech perception in noise than NH listeners. In this scenario, it is possible that the ability to use attention to suppress noise (e.g. the N1-P2_noise_) is not as relevant in CI users. However, this result may have another explanation. Namely, the quality of word recognition in NH listeners may be less variable than in CI users (perhaps due to better preserved input), and therefore variability in NH is more evident in their response to the noise rather than the target word, wherein there is an effective ceiling effect.

Regardless, these data do not argue against the broader process of attentional modulation that was observed by our prior work (Kim et al., 2021). Attentional modulation is likely a bidirectional process, involving both enhancement of a target signal and suppression of a to-be-ignored signal (Noonan et al., 2016). The net result in both the current study and the previous one is still an overall enhancement of internal SNR (i.e., the difference between the cortical response to the signal and the response to the noise) which may underlie better performance. In this light, our data with CI users here thus raises the possibility that different classes of listeners may weight these different aspects of the attentional process differently.

It is worthwhile to consider as to why N1-P2 enhancement is associated with better speech perception outcomes. As with ERPs in other modalities, previous studies have demonstrated that the amplitude of the N1-P2 increases as the physical amplitude of the auditory stimulus increases (e.g. Picton et al., 1970; Ross et al., 1999). Therefore, one interpretation may be that an enhanced N1-P2 complex may simply reflect better sensitivity to auditory events in general in better listeners. This explanation is unlikely to be correct for several reasons. First, we do not see this effect to the noise onset. Thus, variation in the N1-P2 is not due to low level auditory sensitivity. Second, our acoustic hearing thresholds (PTA_low_) provide an estimate of such sensitivity, and we found that neither N1-P2 (to target or to noise) was correlated with this. Finally, even when we controlled for PTA_low_ the relationship between N1P2_target_ and performance was still significant. As a result, a simple peripheral explanation for these results is not well supported here.

It has also been observed previously that the amplitude of the N1-P2 complex increases with a greater saliency of sensory events (Stevenson et al., 2012; Kamal et al., 2021). In this context, we may theorize that an increased cortical response to the target word reflects better separation of speech from the background noise (i.e., increased saliency). Further investigation is required to explore this in greater detail, by understanding the relationship to other factors that reflect the saliency of auditory events, such as measures of spectral and temporal resolution, which were not obtained here.

In normal hearing listeners, Obleser and Kotz (2011) showed that N1 responses to the onsets of a sentence-based task correlated with the comprehension of degraded stimuli. Whilst this supports what we have shown here in terms of ERP amplitudes correlating with speech comprehension, as mentioned previously we did not find such a correlation for the sentence-based AzBio task. We did not examine N1-P2 responses whilst recording behavioral responses to the AzBio task. This leaves open the possibility that one explanation is again that contextual predictive mechanisms involved in a sentence-based task are not relevant for a single word task (such as the CCT) and are therefore not reflected in the ERPs to a task without those mechanisms being recruited.

### Neural generators of the N1-P2 response

The N1-P2_target_ appears to be a useful biomarker of a range of subcortical and cortical processes that are relevant for single word perception in noise and quiet. To understand why, it is important to consider the neural generators of this response and the potential mechanisms underlying this effect.

Previous studies examining the generators of the auditory N1 component of the N1-P2 complex have implicated the auditory cortices in both hemispheres (e.g. Celesia, 1976; Giard et al., 1994, Gander et al., 2010), with some further contribution from frontal and parietal sources. Similarly, the P2 component is thought to arise from both primary and secondary cortices in bilateral Heschl’s gyrus (for a review, see Lightfoot, 2016), although the contributions from more anterior regions may be greater than for the N1 (Ross & Tremblay, 2009), suggestive of a higher-order, secondary area generator that reflects the processing of more complex stimulus features (Howard et al., 2000; Shahin et al., 2005). Meanwhile, MEG work by Lutkenhoner & Steinstrater (1998) suggested that the N1 arises from planum temporale but also may have multiple generators, whilst the P2 was primarily generated from Heschl’s gyrus. Although not explored in detail here, the topographies in our current study supported the idea of the N1-P2 arising from auditory cortex.

In the current data, age was significantly negatively correlated with N1-P2 amplitudes, even when the data were sub-grouped for the purposes of examining performance on the other two tasks. This indicates that N1-P2 amplitudes decreased with age. The data on the effect of age on these components are mixed. One recent study showed that N1 and P2 individually actually increased with age in response to amplitude-modulated stimuli (Irsik et al., 2020). However, other work examining speech stimuli has demonstrated smaller N1 amplitudes and larger P2 amplitudes in older adults (Rufener et al., 2014). Others have also argued that increased cortical response amplitudes in the aging brain reflect an overcompensation to degraded brainstem encoding (Bidelman et al., 2014), not a true change in auditory cortical processing. Resolving this particular debate may require stronger linking functions explaining *why* the N1-P2 is larger or smaller.

Regardless of this, even when age was accounted for, N1-P2 amplitudes still predicted performance on both single word speech perception tasks, highlighting that the prediction from this response reflected neural mechanisms that were not simply a consequence of aging. One candidate for such a mechanism could be degraded temporal processing, resulting in a reduction of neural synchrony and therefore reduced cortical amplitudes (Eggermont, 2015). Under this premise, an inverse perspective would be that better speech perception occurs in some listeners through preservation or restoration of temporal processing via compensatory mechanisms at the level of auditory cortex, even in the presence of mechanisms that otherwise degrade temporal processing (such as aging and hearing loss).

### Limitations

There are several key limitations to the current study. First, as mentioned, we did not obtain any psychophysical estimates of encoding fidelity in our subjects (e.g. spectral ripple or temporal modulation detection). The precision of peripheral encoding is known to contribute significantly to variance in speech perception performance across subjects (Shannon, 1992; Fu, 2002; Jin & Nelson, 2006; Litvak et al., 2007; Won et al., 2007; Luo et al., 2008; Anderson et al., 2011; Anderson et al., 2012). It is as yet unclear whether the N1-P2_target_ may reflect encoding fidelity (in which case one might predict that certain measures of the periphery would mediate the relationship between N1-P2 and speech perception).

Second, we did not account for other factors such as duration of deafness (although a likely close proxy is age of implant, which correlated significantly with age; r = 0.95, *p* < 0.001). Age of onset of hearing loss was not consistently reliably obtained in our sample, in part because many of our patients experienced gradual hearing loss. However, this makes it impossible to reliably compute duration of deafness. In a very large sample of CI patients, Blamey et al. (2013) showed that patients who had experienced profound hearing loss for a longer duration were less likely to show improvements in speech perception in quiet with CI experience. By contrast, results from Jolink et al. (2016) indicated that age at implantation did not significantly affect speech recognition. In this regard it may be useful to see how the N1-P2 is related to duration of deafness, potentially as an index of auditory nerve function.

Finally, it should be noted that although we largely focused on acoustic bases of speech-in-noise performance here, work using functional near-infrared spectroscopy has demonstrated that cross-modal activation of auditory brain regions by visual speech is predictive of speech perception performance and cochlear implant outcomes (Lawler et al., 2015; Anderson et al., 2017; Anderson et al., 2019). Therefore, further research is required to elucidate other factors that mediate speech perception variability across subjects.

Finally, we did not separate N1 and P2 responses here, but rather examined the amplitude of the complex. We applied a non-causal filter to the data, which can result in temporal smearing of a large response (see de Cheveigné & Nelken, 2019) but is useful for helping to remove CI artifacts. Therefore, we did not wish to infer anything from a particular component of the response that could be affected by the amplitude of the other response, and so did not compute these separately. Indeed, we are not elucidating the underlying neural network involved in speech perception here, nor the underlying network involved in speech-in-noise perception (for a review, see Hickok & Poeppel, 2007). Nonetheless, it is plausible that the predictability of the amplitude of the complex is dominated by one particular component. Given that performance on the sentence-in-noise task was not predicted by the amplitude of this response – a task which presumably more heavily recruits higher-order areas to account for factors such as the context within surrounding words (Miller et al., 1951) – it is possible that the N1 component is more dominant in underlying the effects in the single word tasks, reflecting how well the target word can be represented by auditory cortex.

### Summary

Despite some limitations, the current study has identified some of the objective factors underlying differences in SiN performance and captured variance explained by certain relevant predictors. Moreover, these data potentially further highlight that different speech perception tests capture different physiological mechanisms (for example, N1-P2 complex amplitudes did not predict a sentence-based task). Indeed, other studies have highlighted that single word tasks likely require recruitment of different mechanisms to sentence-in-noise tasks (for a discussion, see Geller et al., 2020). Such a point should be considered when implementing and interpreting tests, both in a research environment and within the clinic.

In summary, we have demonstrated here that the amplitude of the N1-P2 complex at a vertex electrode location was the clearest predictor of speech perception in single word identification tasks. Interestingly, in normal hearing users we recently demonstrated that behavioral performance could be predicted by a reduction in amplitude of the N1-P2 response to the noise, indicating that a noise suppression mechanism was in place to improve speech reception (Kim et al., 2021), whereas these current data indicate that cochlear implant listeners’ performance was rather mediated by an increased cortical response to the target stimulus. With further development, this could be easily implemented in the clinic with patients to provide a richer profile of hearing ability above standard audiometry, and could potentially be used as a prognostic indicator, as well as an outcome measure following hearing remediation. Further work would be beneficial to elucidate the precise neural mechanisms underlying this result. Additionally, residual low frequency hearing thresholds were predictive of variability in perceiving sentences in noise and therefore preservation of residual acoustic hearing during cochlear implantation (and beyond) likely benefits patients in common listening environments.

## Author contributions

I.C. designed the experiments. S.K., A.T.S., J.W., Y.N., A.H., and I.C. ran the experiments. J.I.B., P.E.G., J.W., Y.N., C.D., M.H., B.G., B.M., T.D.G., and I.C. analyzed and interpreted data. J.I.B. and I.C. prepared the manuscript, with revisions and suggestions from B.M. and T.D.G.

## Acknowledgments

This work was supported by NIDCD P50 (DC000242 31) awarded to Gantz, Griffiths and McMurray, NIH T32 (DC000040-24) awarded to Hansen, Department of Defense Hearing Restoration and Rehabilitation Program grant awarded to Choi (W81XWH1910637), and DC008089 awarded to McMurray. The authors declare no competing financial interests.

## Supplementary Data

**Figure S1.**
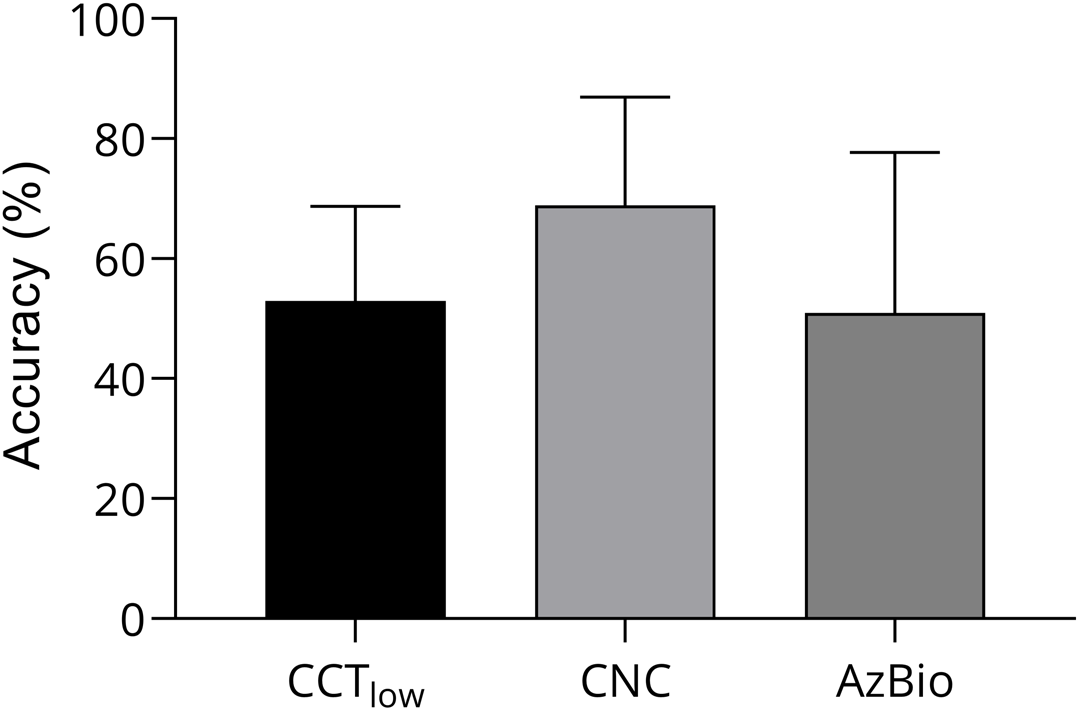
Mean (± S.D.) performance on the three tasks used in the study, across subjects.

**Figure S2.**
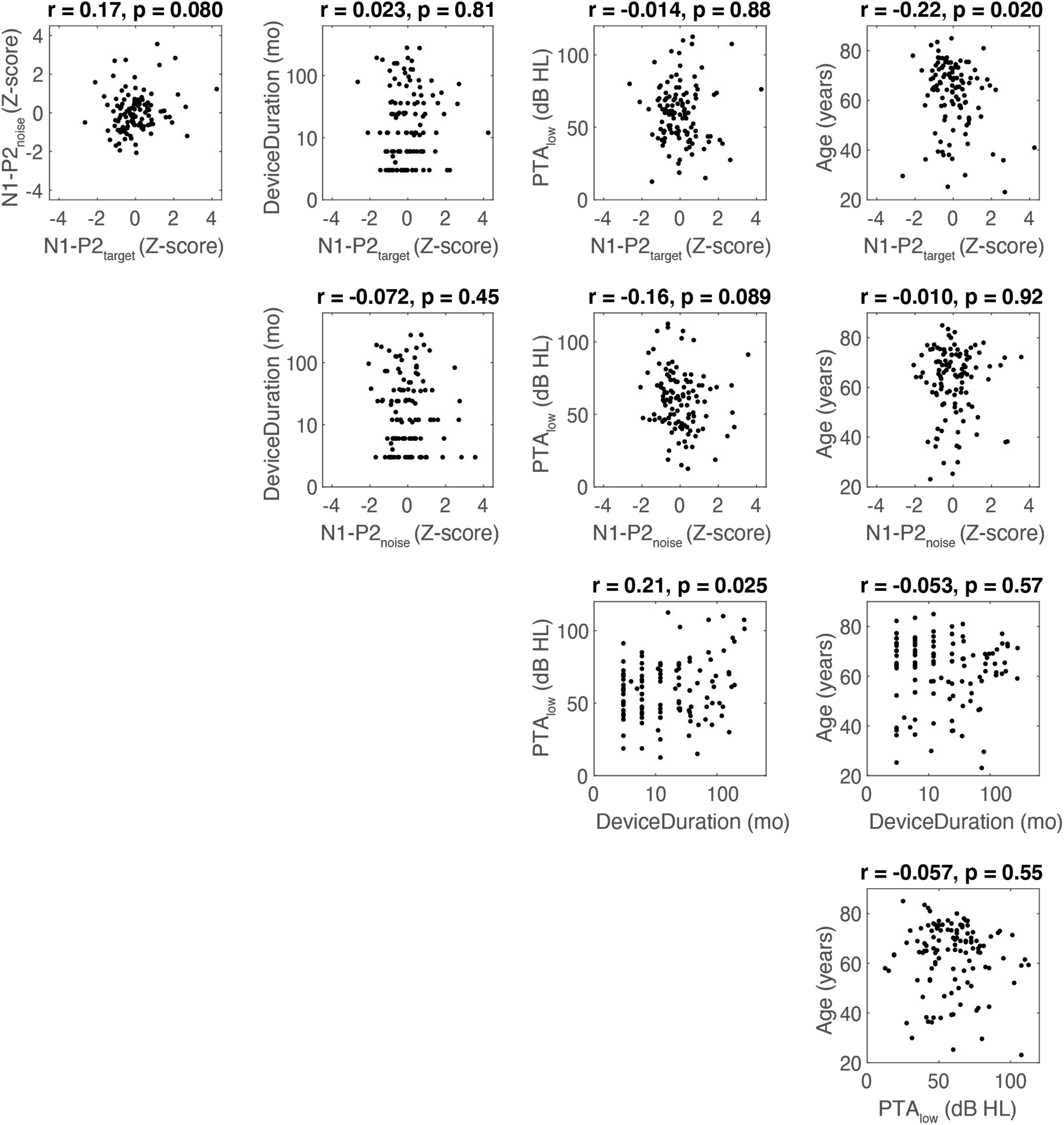
Correlation matrix of predictor variables (n = 114). N1P2_noise_: N1-P2 ERP amplitude at the noise onset. N1P2_target_: N1-P2 ERP amplitude at the target word onset.

**Table S1.** Detailed, anonymized information for all 114 subjects included in the study

**Table S2.** Phonetic balance (Number of items in each phonetic category) of California Consonant Test in each SNR condition

| Target Consonant | SNR: +13 dB | SNR: +7 dB |
| --- | --- | --- |
| Affricate & Fricative | 17 (34%) | 16 (32%) |
| Plosive | 16 (32%) | 17 (34%) |
| Sonorant | 17 (32%) | 17 (34%) |

## References

Anderson, C. A., Wiggins, I. M., Kitterick, P. T., & Hartley, D. E. H. (2017). Adaptive benefit of cross-modal plasticity following cochlear implantation in deaf adults. 114(38), 10256–10261. https://doi.org/10.1073/pnas.1704785114%J Proceedings of the National Academy of Sciences

Anderson, C. A., Wiggins, I. M., Kitterick, P. T., & Hartley, D. E. H. (2019). Pre-operative Brain Imaging Using Functional Near-Infrared Spectroscopy Helps Predict Cochlear Implant Outcome in Deaf Adults. J Assoc Res Otolaryngol, 20(5), 511–528. https://doi.org/10.1007/s10162-019-00729-z

Anderson, E. S., Nelson, D. A., Kreft, H., Nelson, P. B., & Oxenham, A. J. (2011). Comparing spatial tuning curves, spectral ripple resolution, and speech perception in cochlear implant users. J Acoust Soc Am, 130(1), 364–375. https://doi.org/10.1121/1.3589255

Anderson, E. S., Oxenham, A. J., Nelson, P. B., & Nelson, D. A. (2012). Assessing the role of spectral and intensity cues in spectral ripple detection and discrimination in cochlear-implant users. J Acoust Soc Am, 132(6), 3925–3934. https://doi.org/10.1121/1.4763999

Bidelman, G. M., Villafuerte, J. W., Moreno, S., & Alain, C. (2014). Age-related changes in the subcortical-cortical encoding and categorical perception of speech. Neurobiol Aging, 35(11), 2526–2540. https://doi.org/10.1016/j.neurobiolaging.2014.05.006

Blamey, P., Artieres, F., Başkent, D., Bergeron, F., Beynon, A., Burke, E., Dillier, N., Dowell, R., Fraysse, B., Gallégo, S., Govaerts, P. J., Green, K., Huber, A. M., Kleine-Punte, A., Maat, B., Marx, M., Mawman, D., Mosnier, I., O’Connor, A. F., O’Leary, S., Rousset, A., Schauwers, K., Skarzynski, H., Skarzynski, P. H., Sterkers, O., Terranti, A., Truy, E., Van de Heyning, P., Venail, F., Vincent, C., & Lazard, D. S. (2013). Factors affecting auditory performance of postlinguistically deaf adults using cochlear implants: an update with 2251 patients. Audiol Neurootol, 18(1), 36–47. https://doi.org/10.1159/000343189

Brainard, D. H. (1997). The Psychophysics Toolbox. Spat Vis, 10(4), 433–436. https://www.ncbi.nlm.nih.gov/pubmed/9176952

Celesia, G. G. (1976). Organization of auditory cortical areas in man. Brain, 99(3), 403–414. https://doi.org/10.1093/brain/99.3.403

Chang, S. A., Tyler, R. S., Dunn, C. C., Ji, H., Witt, S. A., Gantz, B., & Hansen, M. (2010). Performance over time on adults with simultaneous bilateral cochlear implants. J Am Acad Audiol, 21(1), 35–43. https://doi.org/10.3766/jaaa.21.1.5

de Cheveigné, A., & Nelken, I. (2019). Filters: When, Why, and How (Not) to Use Them. Neuron, 102(2), 280–293. https://doi.org/10.1016/j.neuron.2019.02.039

Debruyne, J. A., Janssen, A. M., & Brokx, J. P. L. (2020). Systematic Review on Late Cochlear Implantation in Early-Deafened Adults and Adolescents: Predictors of Performance. Ear Hear, 41(6), 1431–1441. https://doi.org/10.1097/AUD.0000000000000889

Delorme, A., & Makeig, S. (2004). EEGLAB: an open source toolbox for analysis of single-trial EEG dynamics including independent component analysis. J Neurosci Methods, 134(1), 9–21. https://doi.org/10.1016/j.jneumeth.2003.10.009

Dillon, M. T., Buss, E., Adunka, M. C., King, E. R., Pillsbury, H. C., 3rd, Adunka, O. F., & Buchman, C. A. (2013). Long-term speech perception in elderly cochlear implant users. JAMA Otolaryngol Head Neck Surg, 139(3), 279–283. https://doi.org/10.1001/jamaoto.2013.1814

Dormann, C. F., Elith, J., Bacher, S., Buchmann, C., Carl, G., Carré, G., Marquéz, J. R. G., Gruber, B., Lafourcade, B., Leitão, P. J., Münkemüller, T., McClean, C., Osborne, P. E., Reineking, B., Schröder, B., Skidmore, A. K., Zurell, D., & Lautenbach, S. (2013). Collinearity: a review of methods to deal with it and a simulation study evaluating their performance. 36(1), 27–46. https://doi.org/10.1111/j.1600-0587.2012.07348.x

Du, Y., Buchsbaum, B. R., Grady, C. L., & Alain, C. (2014). Noise differentially impacts phoneme representations in the auditory and speech motor systems. Proc Natl Acad Sci U S A, 111(19), 7126–7131. https://doi.org/10.1073/pnas.1318738111

Dunn, C. C., Perreau, A., Gantz, B., & Tyler, R. S. (2010). Benefits of localization and speech perception with multiple noise sources in listeners with a short-electrode cochlear implant. J Am Acad Audiol, 21(1), 44–51. https://doi.org/10.3766/jaaa.21.1.6

Eggermont, J. J. (2015). Auditory Temporal Processing and its Disorders. Oxford University Press. https://doi.org/10.1093/acprof:oso/9780198719090.001.0001

Finke, M., Buchner, A., Ruigendijk, E., Meyer, M., & Sandmann, P. (2016). On the relationship between auditory cognition and speech intelligibility in cochlear implant users: An ERP study. Neuropsychologia, 87, 169–181. https://doi.org/10.1016/j.neuropsychologia.2016.05.019

Fu, Q. J. (2002). Temporal processing and speech recognition in cochlear implant users. Neuroreport, 13(13), 1635–1639. https://doi.org/10.1097/00001756-200209160-00013

Fullgrabe, C., Moore, B. C., & Stone, M. A. (2014). Age-group differences in speech identification despite matched audiometrically normal hearing: contributions from auditory temporal processing and cognition. Front Aging Neurosci, 6, 347. https://doi.org/10.3389/fnagi.2014.00347

Gander, P. E., Bosnyak, D. J., & Roberts, L. E. (2010). Acoustic experience but not attention modifies neural population phase expressed in human primary auditory cortex. Hear Res, 269(1-2), 81–94. https://doi.org/10.1016/j.heares.2010.07.001

Gantz, B. J., Hansen, M. R., Turner, C. W., Oleson, J. J., Reiss, L. A., & Parkinson, A. J. (2009). Hybrid 10 clinical trial: preliminary results. Audiol Neurootol, 14 Suppl 1, 32–38. https://doi.org/10.1159/000206493

Gantz, B. J., Turner, C., Gfeller, K. E., & Lowder, M. W. (2005). Preservation of hearing in cochlear implant surgery: advantages of combined electrical and acoustical speech processing. Laryngoscope, 115(5), 796–802. https://doi.org/10.1097/01.MLG.0000157695.07536.D2

Gantz, B. J., Woodworth, G. G., Knutson, J. F., Abbas, P. J., & Tyler, R. S. (1993). Multivariate predictors of audiological success with multichannel cochlear implants. Ann Otol Rhinol Laryngol, 102(12), 909–916. https://doi.org/10.1177/000348949310201201

Gay, J. D., Voytenko, S. V., Galazyuk, A. V., & Rosen, M. J. (2014). Developmental hearing loss impairs signal detection in noise: putative central mechanisms. Front Syst Neurosci, 8, 162. https://doi.org/10.3389/fnsys.2014.00162

Geller, J., Holmes, A., Schwalje, A., Berger, J. I., Gander, P. E., Choi, I., & McMurray, B. (2020). Validating the Iowa Test of Consonant Perception. PsyArXiv. https://doi.org/10.31234/osf.io/wxd93

Getz, L. M., & Toscano, J. C. (2020). The time-course of speech perception revealed by temporally-sensitive neural measures. Wiley Interdisciplinary Reviews-Cognitive Science. https://doi.org/ARTN e1541 10.1002/wcs.1541

Giard, M. H., Perrin, F., Echallier, J. F., Thevenet, M., Froment, J. C., & Pernier, J. (1994). Dissociation of temporal and frontal components in the human auditory N1 wave: a scalp current density and dipole model analysis. Electroencephalogr Clin Neurophysiol, 92(3), 238–252. https://doi.org/10.1016/0168-5597(94)90067-1

Gifford, R. H., Dorman, M. F., Skarzynski, H., Lorens, A., Polak, M., Driscoll, C. L., Roland, P., & Buchman, C. A. (2013). Cochlear implantation with hearing preservation yields significant benefit for speech recognition in complex listening environments. Ear Hear, 34(4), 413–425. https://doi.org/10.1097/AUD.0b013e31827e8163

Green, K. M., Bhatt, Y., Mawman, D. J., O’Driscoll, M. P., Saeed, S. R., Ramsden, R. T., & Green, M. W. (2007). Predictors of audiological outcome following cochlear implantation in adults. Cochlear Implants Int, 8(1), 1–11. https://doi.org/10.1179/cim.2007.8.1.1

Groenen, P. A., Beynon, A. J., Snik, A. F., & van den Broek, P. (2001). Speech-evoked cortical potentials and speech recognition in cochlear implant users. Scand Audiol, 30(1), 31–40. https://doi.org/10.1080/010503901750069554

Groenen, P. A., Makhdoum, M., van den Brink, J. L., Stollman, M. H., Snik, A. F., & van den Broek, P. (1996). The relation between electric auditory brain stem and cognitive responses and speech perception in cochlear implant users. Acta Otolaryngol, 116(6), 785–790. https://doi.org/10.3109/00016489609137926

Guest, H., Munro, K. J., Prendergast, G., Millman, R. E., & Plack, C. J. (2018). Impaired speech perception in noise with a normal audiogram: No evidence for cochlear synaptopathy and no relation to lifetime noise exposure. Hear Res, 364, 142–151. https://doi.org/10.1016/j.heares.2018.03.008

Harris, R. J. (2001). A Primer of Multivariate Statistics (3rd ed.).. Psychology Press. https://doi.org/10.4324/9781410600455

Hickok, G., & Poeppel, D. (2007). The cortical organization of speech processing. Nat Rev Neurosci, 8(5), 393–402. https://doi.org/10.1038/nrn2113

Howard, M. A., Volkov, I. O., Mirsky, R., Garell, P. C., Noh, M. D., Granner, M., Damasio, H., Steinschneider, M., Reale, R. A., Hind, J. E., & Brugge, J. F. (2000). Auditory cortex on the human posterior superior temporal gyrus. J Comp Neurol, 416(1), 79–92. https://doi.org/10.1002/(sici)1096-9861(20000103)416:1<79::aid-cne6>3.0.co;2-2

Irsik, V. C., Almanaseer, A., Johnsrude, I. S., & Herrmann, B. (2020). Cortical Responses to the Amplitude Envelopes of Sounds Change with Age. 2020.2010.2023.352880. https://doi.org/10.1101/2020.10.23.352880%J bioRxiv

Jin, S. H., & Nelson, P. B. (2006). Speech perception in gated noise: the effects of temporal resolution. J Acoust Soc Am, 119(5 Pt 1), 3097–3108. https://doi.org/10.1121/1.2188688

Jolink, C., Helleman, H. W., van Spronsen, E., Ebbens, F. A., Ravesloot, M. J., & Dreschler, W. A. (2016). The long-term results of speech perception in elderly cochlear implant users. Cochlear Implants Int, 17(3), 146–150. https://doi.org/10.1080/14670100.2016.1162383

Kamal, F., Morrison, C., Campbell, K., & Taler, V. (2021). Event-related potential evidence that very slowly presented auditory stimuli are passively processed differently in younger and older adults. Neurobiology of Aging. https://doi.org/10.1016/j.neurobiolaging.2021.02.014

Kim, S., Schwalje, A. T., Liu, A. S., Gander, P. E., McMurray, B., Griffiths, T. D., & Choi, I. (2021). Pre- and post-target cortical processes predict speech-in-noise performance. Neuroimage, 228, 117699. https://doi.org/10.1016/j.neuroimage.2020.117699

Kitterick, P. T., & Lucas, L. (2016). Predicting speech perception outcomes following cochlear implantation in adults with unilateral deafness or highly asymmetric hearing loss. Cochlear Implants Int, 17 Suppl 1, 51–54. https://doi.org/10.1080/14670100.2016.1155806

Lawler, C. A., Wiggins, I. M., Dewey, R. S., & Hartley, D. E. (2015). The use of functional near-infrared spectroscopy for measuring cortical reorganisation in cochlear implant users: a possible predictor of variable speech outcomes? Cochlear Implants Int, 16 Suppl 1, S30–32. https://doi.org/10.1179/1467010014z.000000000230

Lehiste, I., & Peterson, G. E. (1959). Linguistic Considerations in the Study of Speech Intelligibility. 31(3), 280–286. https://doi.org/10.1121/1.1907713

Liberman, M. C., Epstein, M. J., Cleveland, S. S., Wang, H., & Maison, S. F. (2016). Toward a Differential Diagnosis of Hidden Hearing Loss in Humans. PLoS One, 11(9), e0162726. https://doi.org/10.1371/journal.pone.0162726

Lightfoot, G. (2016). Summary of the N1-P2 Cortical Auditory Evoked Potential to Estimate the Auditory Threshold in Adults. Semin Hear, 37(1), 1–8. https://doi.org/10.1055/s-0035-1570334

Litvak, L. M., Spahr, A. J., Saoji, A. A., & Fridman, G. Y. (2007). Relationship between perception of spectral ripple and speech recognition in cochlear implant and vocoder listeners. 122(2), 982–991. https://doi.org/10.1121/1.2749413

Luo, X., Fu, Q. J., Wei, C. G., & Cao, K. L. (2008). Speech recognition and temporal amplitude modulation processing by Mandarin-speaking cochlear implant users. Ear Hear, 29(6), 957–970. https://doi.org/10.1097/AUD.0b013e3181888f61

Lutkenhoner, B., & Steinstrater, O. (1998). High-precision neuromagnetic study of the functional organization of the human auditory cortex. Audiol Neurootol, 3(2-3), 191–213. https://doi.org/10.1159/000013790

Makhdoum, M. J. A., Hinderink, J. B., Snik, A. F. M., Groenen, P., & Van Den Broek, P. (1998). Can event-related potentials be evoked by extra-cochlear stimulation and used for selection purposes in cochlear implantation?, 23(5), 432–438. https://doi.org/10.1046/j.1365-2273.1998.00168.x

Maris, E., & Oostenveld, R. (2007). Nonparametric statistical testing of EEG- and MEG-data. J Neurosci Methods, 164(1), 177–190. https://doi.org/10.1016/j.jneumeth.2007.03.024

McCormack, A., & Fortnum, H. (2013). Why do people fitted with hearing aids not wear them? Int J Audiol, 52(5), 360–368. https://doi.org/10.3109/14992027.2013.769066

McMurray, B., Munson, C., & Tomblin, J. B. (2014). Individual differences in language ability are related to variation in word recognition, not speech perception: evidence from eye movements. J Speech Lang Hear Res, 57(4), 1344–1362. https://doi.org/10.1044/2014_JSLHR-L-13-0196

Micco, A. G., Kraus, N., Koch, D. B., McGee, T. J., Carrell, T. D., Sharma, A., Nicol, T., & Wiet, R. J. (1995). Speech-evoked cognitive P300 potentials in cochlear implant recipients. Am J Otol, 16(4), 514–520. https://www.ncbi.nlm.nih.gov/pubmed/8588653

Miller, G. A., Heise, G. A., & Lichten, W. (1951). The intelligibility of speech as a function of the context of the test materials. Journal of Experimental Psychology, 41(5), 329–335. https://doi.org/10.1037/h0062491

Noonan, M. P., Adamian, N., Pike, A., Printzlau, F., Crittenden, B. M., & Stokes, M. G. (2016). Distinct Mechanisms for Distractor Suppression and Target Facilitation. J Neurosci, 36(6), 1797–1807. https://doi.org/10.1523/JNEUROSCI.2133-15.2016

Obleser, J., & Kotz, S. A. (2011). Multiple brain signatures of integration in the comprehension of degraded speech. Neuroimage, 55(2), 713–723. https://doi.org/10.1016/j.neuroimage.2010.12.020

Owens, E., & Schubert, E. D. (1977). Development of the California Consonant Test. J Speech Hear Res, 20(3), 463–474. https://doi.org/10.1044/jshr.2003.463

Pelli, D. G. (1997). The VideoToolbox software for visual psychophysics: transforming numbers into movies. Spat Vis, 10(4), 437–442. https://www.ncbi.nlm.nih.gov/pubmed/9176953

Picton, T. W., Goodman, W. S., & Bryce, D. P. (1970). Amplitude of Evoked Responses to Tones of High Intensity. Acta Oto-Laryngologica, 70(2), 77–82. https://doi.org/10.3109/00016487009181862

Purdy, S. C., & Kelly, A. S. (2016). Change in Speech Perception and Auditory Evoked Potentials over Time after Unilateral Cochlear Implantation in Postlingually Deaf Adults. Semin Hear, 37(1), 62–73. https://doi.org/10.1055/s-0035-1570329

Ross, B., Lütkenhöner, B., Pantev, C., & Hoke, M. (1999). Frequency-Specific Threshold Determination with the CERAgram Method: Basic Principle and Retrospective Evaluation of Data. Audiology and Neurotology, 4(1), 12–27. https://doi.org/10.1159/000013816

Ross, B., & Tremblay, K. (2009). Stimulus experience modifies auditory neuromagnetic responses in young and older listeners. Hear Res, 248(1-2), 48–59. https://doi.org/10.1016/j.heares.2008.11.012

Rubinstein, J. T., Parkinson, W. S., Tyler, R. S., & Gantz, B. J. (1999). Residual speech recognition and cochlear implant performance: effects of implantation criteria. Am J Otol, 20(4), 445–452. https://www.ncbi.nlm.nih.gov/pubmed/10431885

Rufener, K. S., Liem, F., & Meyer, M. (2014). Age-related differences in auditory evoked potentials as a function of task modulation during speech-nonspeech processing. Brain Behav, 4(1), 21–28. https://doi.org/10.1002/brb3.188

Saiz-Alia, M., Forte, A. E., & Reichenbach, T. (2019). Individual differences in the attentional modulation of the human auditory brainstem response to speech inform on speech-in-noise deficits. Sci Rep, 9(1), 14131. https://doi.org/10.1038/s41598-019-50773-1

Sarrett, M. E., McMurray, B., & Kapnoula, E. C. (2020). Dynamic EEG analysis during language comprehension reveals interactive cascades between perceptual processing and sentential expectations. Brain and Language, 211. https://doi.org/ARTN 104875 10.1016/j.bandl.2020.104875

Shahin, A., Roberts, L. E., Pantev, C., Trainor, L. J., & Ross, B. (2005). Modulation of P2 auditory-evoked responses by the spectral complexity of musical sounds. Neuroreport, 16(16), 1781–1785. https://doi.org/10.1097/01.wnr.0000185017.29316.63

Shannon, R. V. (1992). Temporal modulation transfer functions in patients with cochlear implants. J Acoust Soc Am, 91(4 Pt 1), 2156–2164. https://doi.org/10.1121/1.403807

Spahr, A. J., Dorman, M. F., Litvak, L. M., Van Wie, S., Gifford, R. H., Loizou, P. C., Loiselle, L. M., Oakes, T., & Cook, S. (2012). Development and validation of the AzBio sentence lists. Ear Hear, 33(1), 112–117. https://doi.org/10.1097/AUD.0b013e31822c2549

Stevenson, R. A., Bushmakin, M., Kim, S., Wallace, M. T., Puce, A., & James, T. W. (2012). Inverse effectiveness and multisensory interactions in visual event-related potentials with audiovisual speech. Brain Topogr, 25(3), 308–326. https://doi.org/10.1007/s10548-012-0220-7

Taylor, B. (2003). Speech-in-noise tests: How and why to include them in your basic test battery. The Hearing Journal, 56(1), 40–42. https://doi.org/10.1097/01.HJ.0000293000.76300.ff

Tremblay, K., Kraus, N., McGee, T., Ponton, C., & Otis, B. (2001). Central auditory plasticity: changes in the N1-P2 complex after speech-sound training. Ear Hear, 22(2), 79–90. https://doi.org/10.1097/00003446-200104000-00001

Turner, C. W., Gantz, B. J., Vidal, C., Behrens, A., & Henry, B. A. (2004). Speech recognition in noise for cochlear implant listeners: benefits of residual acoustic hearing. J Acoust Soc Am, 115(4), 1729–1735. https://doi.org/10.1121/1.1687425

Tyler, R. S., Parkinson, A. J., Woodworth, G. G., Lowder, M. W., & Gantz, B. J. (1997). Performance over time of adult patients using the Ineraid or nucleus cochlear implant. J Acoust Soc Am, 102(1), 508–522. https://doi.org/10.1121/1.419724

Won, J. H., Drennan, W. R., & Rubinstein, J. T. (2007). Spectral-ripple resolution correlates with speech reception in noise in cochlear implant users. J Assoc Res Otolaryngol, 8(3), 384–392. https://doi.org/10.1007/s10162-007-0085-8

Wong, P. C., Uppunda, A. K., Parrish, T. B., & Dhar, S. (2008). Cortical mechanisms of speech perception in noise. J Speech Lang Hear Res, 51(4), 1026–1041. https://doi.org/10.1044/1092-4388(2008/075)

